# Deep Local Analysis estimates effects of mutations on protein-protein interactions

**DOI:** 10.1101/2022.10.09.511484

**Authors:** Yasser Mohseni Behbahani, Elodie Laine, Alessandra Carbone

## Abstract

The spectacular advances in protein and protein complex structure prediction hold promises for the reconstruction of interactomes at large scale at the residue resolution. Beyond determining the 3D arrangement of interacting partners, modeling approaches should be able to sense the impact of sequence variations such as point mutations on the strength of the association. In this work, we report on DLA-mutation, a novel and efficient deep learning framework for accurately predicting mutation-induced binding affinity changes. It relies on a 3D-invariant description of local 3D environments at protein interfaces and leverages the large amounts of available protein complex structures through self-supervised learning. It combines the learnt representations with evolutionary information, and a description of interface structural regions, in a siamese architecture. DLA-mutation achieves a Pearson correlation coefficient of 0.81 on a large collection of more than 2000 mutations, and its generalization capability to unseen complexes is higher than state-of-the-art methods.

## 1 Introduction

Disease-related mutations are preferentially localized on protein interaction surfaces compared to other regions of the protein [37, 10, 9]. These mutations affect the propensity and strength with which two proteins interact. Several computational methods have been proposed for estimating mutation-induced binding affinity changes (ΔΔ*G*_*bind*_) by exploiting protein sequence and structural information [38, 15, 29, 31, 32, 36, 40, 23]. In particular, ΔΔ*G*_*bind*_ was found correlated with atomic-distance patterns surrounding the residue subject to the mutation [28, 29, 31, 32]. This correlation emphasises the importance of accounting for local geometrical and physico-chemical environments around the mutation site. Recent methods have been leveraging powerful deep learning techniques to extract abstract representations of the data. For instance, TopNetTree [36] applies convolutional neural networks (CNN) and gradient-boosting trees on a set of pre-computed features representing geometric, topological and contact patterns within the neighbourhood of the mutation site. GraphPPI [23] further obviates the need for feature engineering by using a graph neural network (GNN) that automatically extracts graphical features from an input 3D structure. The GNN is pre-trained on a large body of protein complex structures through self-supervised learning. Self-supervised representation learning has proven successful for predicting various protein structural and functional properties in the context of protein language models [30, 12, 3], for fixed-backbone protein design [1, 17, 8], and for protein stability predictions [5, 39].

Here, we report on *Deep Local Analysis(DLA)-Mutation*, a novel and efficient deep learning-based approach predicting mutation-induced binding affinity changes from local 3D environments around the mutation site (**Fig. 1**). It builds on the DLA framework we previously introduced for assessing the quality of protein complex conformations [25]. DLA applies 3D convolutions to locally oriented residue-centred cubes encapsulating atomic-resolution geometrical and physico-chemical information [27] (**Fig. 1A**). In this work, we expanded this framework by combining self-supervised representation learning of 3D local interfacial environments (**Fig. 1B**) with supervised learning of ΔΔ*G*_*bind*_ exploiting both structural and evolutionary information (**Fig. 1C**). DLA-Mutation only takes as input two cubes, corresponding to the environments around the wild-type and mutated residues, respectively, and directly estimates ΔΔ*G*_*bind*_. It effectively bypasses the estimation of the binding affinities Δ*G*_*bind*_ of the wild-type and mutated complexes, thus avoiding the accumulation of errors on these quantities and allowing for large-scale applications.

**Figure 1:**
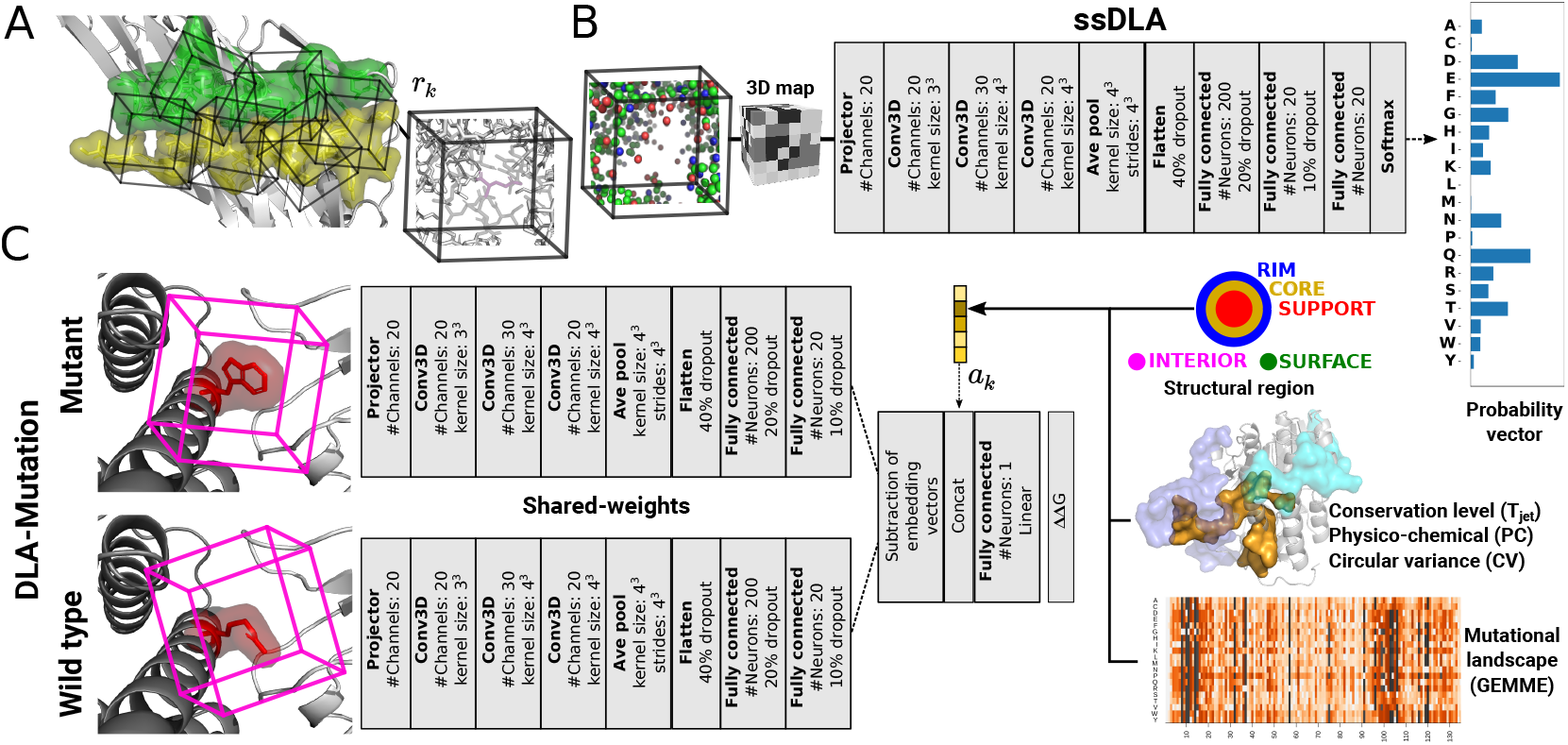
The architectures of DLA framework. **A**. A representation of a protein interface (green and yellow residues from each partner) as an ensemble of cubes (*I*_*C*_). Each cube (*r*_*k*_ *∈ I*_*C*_) is centered and oriented around an interfacial residue. The central residue is masked and the cube contains only those atoms that belong to the local environment. **B**. The pre-trained architecture (ssDLA) is described in detail. It requires a self-supervised training, where the task is to predict the amino acid type from its environment. **C**. Siamese architecture (DLA-Mutation) to predict the changes of binding affinity upon point mutations. Two parallel branches extract descriptive features from wild-type and mutant amino acids. Auxiliary features can be concatenated to the subtraction between the embeddings from two branches to improve the performance.

## 2 Methods

### 2.1 Protein–protein interface representation

We represent a protein-protein interface as a set of locally oriented cubic volumetric maps centered around each interfacial residue (**Fig. 1**A) (See Supplementary Information for details on building the cubic volumetric map). We define interfacial residues as those displaying a change in solvent accessibility between the free (isolated) protein and the complex [21]. We used NACCESS [18] with a probe radius of 1.4Å to compute residue solvent accessibility.

#### Masking procedure

The common practice when applying self-supervised learning to protein sequences is to reconstruct some masked or the next amino acid(s), given their sequence context. This task proved successful for natural language processing [11] before being transferred to proteins. We employed a similar strategy here, by training DLA to recognize which amino acid would fit in a given local 3D environment extracted from a protein-protein interface. Our aim in doing so is to capture intrinsic patterns underlying the atomic arrangements found in local interfacial regions. Formally, the machine predicts the probability *P*(*y*|*env*) of the amino acid type *y*, for *y* ∈ {*A, C, D*, …, *W, Y*}, conditioned on the interfacial local chemical environment *env* given as input. In practice, we process the input cube before giving it to DLA by masking a sphere of radius *r*_*c*_Å centered on an atom from the central residue. Masking a fixed volume prevents introducing amino acid-specific biases. We experimented with different values of *r*_*c*_ (3 and 5Å) and different choices for the atom (*C*_*α*_, *C*_*β*_, random). We found that a sphere of radius of 5Å with a randomly chosen center yielded both good performance and expressive embedding vectors.

#### Auxiliary features

For predicting ΔΔ*G*_*bind*_, we combined the embeddings of the volumetric maps with five pre-computed auxiliary features (**Fig. 1**C), among which four describe the wild-type residue: (i) a one-hot vector encoding its structural region, either the protein interior (INT), the surface (SUR), or, if it is part of the interface, the support (S or SUP), the core (C or COR), or the rim (R or RIM) [21], (ii) its evolutionary conservation level *T*_*JET*_ (a float value) computed by JET [13], (iii) its physico-chemical propensity (PC, a float value) to be found at interfaces [26], and (iv) its circular variance (CV, a float value) [24, 6] reflecting the extent to which it is buried inside the protein. The fifth feature is a numerical score (a float value) estimating the functional impact of the mutation on the monomeric protein. The score is computed by GEMME [20] from a multiple sequence alignment.

### 2.2 DLA architectures

We used the same core architecture for the self-supervised representation learning (**Fig. 1B**) and for the supervised prediction of ΔΔ*G*_*bind*_ (**Fig. 1C**). It comprises a projector, three 3D convolutional layers, an average pooling layer, and a fully connected subnetwork. The projector maps the feature vector of each voxel from the input cube into a vector of size 20. Each convolutional layer is followed by a batch normalization layer. To avoid overfitting, we applied 40%, 20%, and 10% dropout regularization to the input, the first and the second layers of the fully connected subnetwork. The fully-connected subnetwork of the *self supervised-DLA* (*ssDLA*) architecture comprises three successive layers of size 200, 20, and 20 (**Fig. 1B**). The last activation function (Softmax) outputs a probability vector of size 20 representing the 20 amino acids. ssDLA’s loss function is the categorical cross-entropy. The ΔΔ*G*_*bind*_ predictor, DLA-Mutation, processes two input cubes in parallel, corresponding to the wild-type and mutated residues, thanks to a Siamese architecture (**Fig. 1C**). Within each branch, the average pooling layer is followed by two fully connected layers of size 200 and 20. The branches are then merged by subtracting the computed embeddings, and the auxiliary features are concatenated to the resulting vector. The last layer is fully-connected, with one output and linear activation function. The loss function is the mean squared error.

### 2.3 Databases

#### Experimental values for ΔΔ*G*_*bind*_

We used SKEMPI v2.0 [19], the most complete source for experimentally measured binding affinities of wild-type and mutated protein complexes. It reports measurements for over 7 000 point mutations coming from 345 protein complexes, including antibody-antigen (AB/AG) and protease-inhibitor (Pr/PI) assemblies, and assemblies formed between major histocompatibility complex proteins and T-cell receptors (pMHC-TCR). We selected a subset of 2 003 single-point mutations associated with 142 complexes. We call this subset *S2003*.

#### Protein-protein complex 3D structures

We created two databases of protein complex structures, namely *PDBInter* and *S2003-3D*, for training and validation purposes. PDBInter contains 5 055 complex experimental structures curated from the PDB [4]. They do not share any family level similarity between them nor with the 142 complexes from S2003, according to the SCOPe hierarchy [14, 7]. S2003-3D contains 3D models of the wild-type and mutated complexes from S2003. They were generated using the “backrub” protocol implemented in Rosetta [34]

See Supplementary Information for more details about the mutation selection process and the generation of the structural databases.

### 2.4 Training and evaluation protocols

#### Training and validation of ssDLA

We divided the PDBInter database into train and validation sets at the level of complexes. We generated 247 662 interfacial cubes from the training set and 34 174 from the validation set. In both sets, we observed some differences in the frequencies of occurrence of the different amino acids. Leucine is the most frequent one, while cysteine is the rarest (**Fig. SI 1**). To compensate for such imbalance and with the aim of penalizing more the errors made for the less frequent amino acids, we assigned a weight to the loss of each amino acid type that is inversely proportional to its frequency of occurrence. We trained ssDLA for 50 epochs (**Fig. SI 2**).

#### Training and validation of DLA-Mutation

We trained DLA-Mutation by fine-tuning the weights of ssDLA and integrating all the auxiliary features. Machine learning approaches for predicting mutation-induced ΔΔ*G*_*bind*_ typically consider each sample independently when splitting the data between train and test sets [15, 31, 32, 36, 40]. However, this assumption is not valid since several samples may correspond to different mutations taking place in the same complex or even the same position of a complex. Here, we assessed the two types of split, namely mutation-based, where all samples are treated independently, and complex-based, where we guaranteed that no complex was shared between the train and test sets. We performed 10-fold cross validation only with the mutation-based splitting procedure. For the complex-based one, we hold out 32 complexes displaying 391 mutations for the testing phase, and trained DLA-Mutation on the rest of the dataset. For comparison with other predictors we selected 112 mutations from 17 complexes as the test set. This set is the intersection between S2003 and the benchmark set for which the results of the other methods were reported in [15]. We retrained the model using 945 mutations from S2003 with complexes sharing less than 30% sequence identity with those from test set.

## 3 Results

### 3.1 Inferring interfacial amino acid types from 3D local environments

We first assessed the ability of ssDLA to recover the identity of a masked amino acid from its local environment (**Fig. 2A**). ssDLA successfully and consistently recognises the amino acids containing an aromatic ring, namely F, Y, W, H, and most of the charged and polar ones, namely E, K, R, and to a lesser extent Q and D, as well as methionine (M), cysteine (C), glycine (G), and proline (P), whatever their structural region. By contrast, the location of alanine (A), isoleucine (I) and leucine (L) influences their detection. While they are ranked in the top 3 in the support and the core, they are almost never recognised in the rim. Inversely, the polar asparagine (N) is recognised when located in the rim or the core, but not the support. The model often confuses the hydroxyl-containing serine (S) and threonine (T) on the one hand, and the hydrophobic I and L on the other hand. Overall, it tends to over-populate the rim with aspartate (D). These tendencies differ from those reported previously for a similar task and data representation [1]. In particular, the model from [1] identifies glycine and proline with very high success and tends to confuse F, Y and W. Such differences may be explained by the fact that the authors masked the side chain of the central residue, instead of a constant volume. This choice may encourage the model to put more importance on the size and the shape of the missing part in making its prediction, than on its physico-chemical environment. Let us also stress that the model reported in [1] is trained on monomeric proteins.

**Figure 2:**
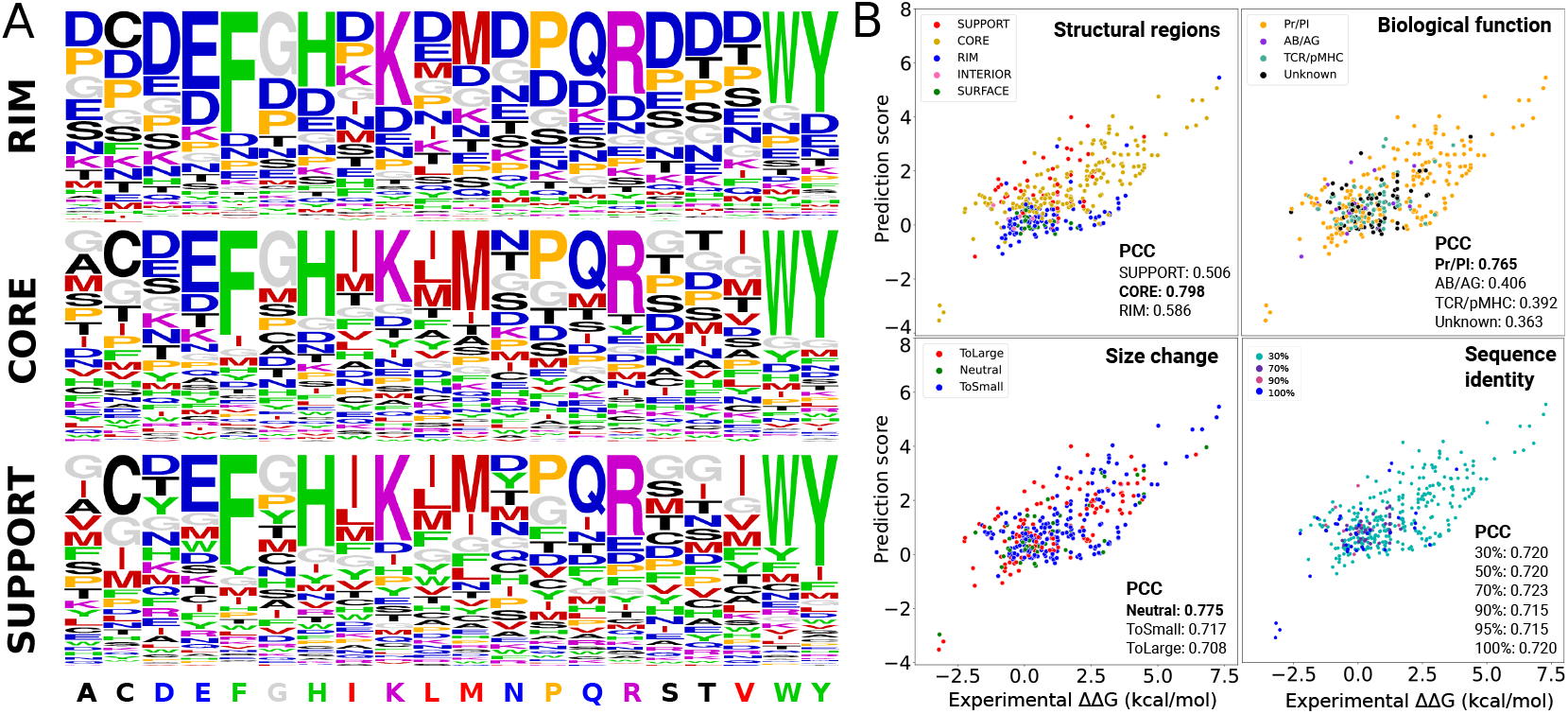
Predictive performance of DLA-Mutation. **A**. Recovery of the central amino acid by ssDLA in the self-supervised representation task, using the validation set of PDBInter. The three logos represent the propensities of each amino acid to be predicted (having maximum score in the output layer), depending on the true amino acid (x-axis) and on its structural region (see *Methods*). The colors indicate geometrical and physico-chemical properties (**Table SI 1**). **B**. Performance in predicting ΔΔ*G*_*bind*_ on the testing set of 391 mutations from 32 unseen protein complexes classified by different characteristics. The overall performance is PCC=0.72. The dots are colored with respect to the structural region of the mutant residue (top left), the biological function of the protein complex (top right), the change in the size between wild-type and mutant residue (bottom left) and the minimum sequence identity shared with any training complex.

### 3.2 DLA-Mutation outperforms state-of-the-art ΔΔG predictors

We used Pearson correlation coefficient (PCC) and root mean squared error (RMSE) as the evaluation metrics. DLA-Mutation achieved a PCC = 0.812 in the 10-fold mutation-based cross validation procedure (**Fig. SI 5**). We further evaluated DLA-Mutation using the complex-based split with the test set of 391 mutations. Combined with the auxiliary features, DLA-Mutation reached a PCC = 0.720 in ΔΔG prediction. We also investigated the influence of the mutated residue’s structural region (support, core, rim, interior or surface), the complex biological function, or the amino acid size change upon mutation (**Fig. 2**). 89% of the mutations are located on the interface with the majority of them belonging to the core. DLA-Mutation performs better on the core residues (PCC=0.798) than those from rim (PCC=0.586) and support (PCC=0.506). The mutations in the support display the highest variability. The majority of mutations happen on the interface of protease-inhibitor assemblies. DLA-Mutation performs well for this subset with PCC=0.765. The performance drops on the other class down to 0.3-0.4. This might be explained by the low diversity of substitutions in these classes, with the overwhelming majority (> 84%) of the mutations being substitutions to alanine. Although the predictions of DLA-Mutation are robust to the change of the amino acid size, the correlation is better for the neutral group (PCC=0.775) (See Supplementary Information for details). Our approach is robust to sequence identity overlap between train and test sets (**Fig. 2**). The complexes of the majority of the mutations share less than 30 % sequence identity with the training set. Finally, we compared the performance of DLA-Mutation with those of BindProfX, FoldX, iSEE and mCSM on 17 complexes displaying 112 mutations (**Fig. 3**). For this subset, DLA-Mutation reached PCC=0.48, outperforming the other approaches.

**Figure 3:**
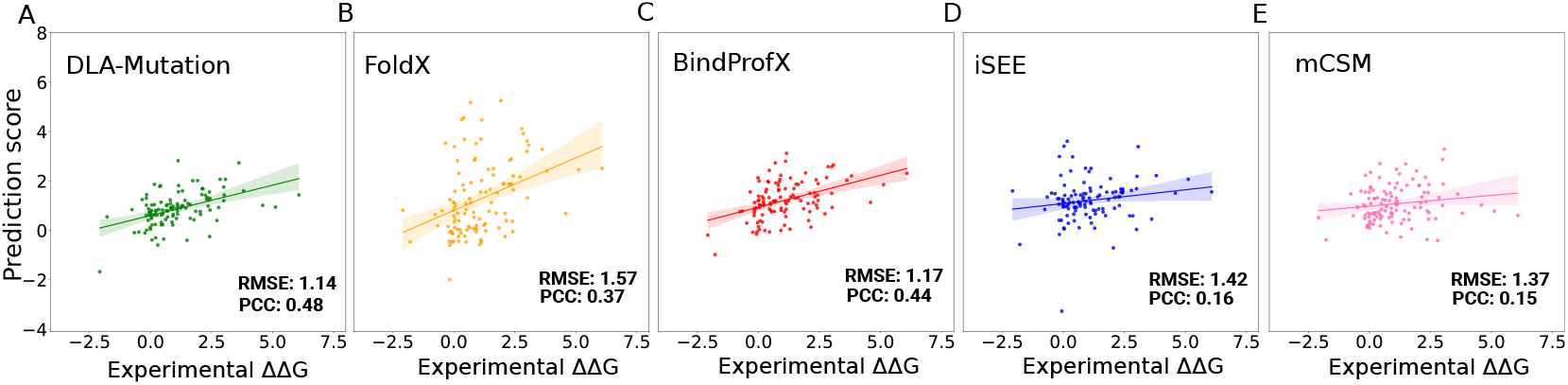
A comparison between DLA-Mutation and other approaches in the prediction of ΔΔ*G*. Performance on 112 mutations associated to 17 complexes for DLA-Mutation (**A**), FoldX (**B**), BindProfX (**C**), iSEE (**D**) and mCSM (**E**).

## 4 Discussion

We have proposed a deep learning-based method for assessing the impact of mutations on protein-protein binding affinity. It derives and contrasts representations of the local geometrical and physico-chemical environments around the mutation site in the wild-type and mutated forms with a Siamese architecture. The representations are enriched with evolutionary information coming from sequences related to the protein bearing the mutation. Beyond improving the prediction of ΔΔ*G*_*bind*_ over the state of the art, it would be interesting to investigate what the representations learnt during the self-supervised step can tell us about the specificity of interfacial environments and the functions of protein interactions. Future developments will also aim at systematically scanning protein complex interfaces on a proteome-wide scale.

## Acknowledgements

We thank the Institute for Development and Resources in Intensive Scientific Computing (IDRIS-CNRS) for giving us access to their Jean Zay supercomputer.

## Supplementary Information

### Building the cubic volumetric map

To build the cubic volumetric map, the atomic coordinates of the input structure are first transformed to a density function [27]. The density *d* at a point 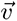 is computed as

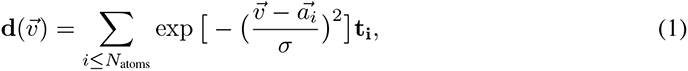

Where 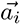 is the position of the *ith* atom, σ is the width of the Gaussian kernel set to 1Å, and *t*_*i*_ is a vector of 167 channels that correspond to residue-specific atom types (O, C, N and S). The hydrogen atoms are discarded. Then, the density is projected on a 3D grid comprising 24 × 24 × 24 voxels of side 0.8Å. The map is oriented by defining a local frame based on the common chemical scaffold of amino acid residues in proteins [27]. More precisely, for the *nth* residue, the 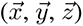 directions and the origin of the cube are defined by the position of the atom N_*n*_, and the directions of C_*n*−1_ and Cα_*n*_ with respect to N_*n*_. The X-axis is parallel to the vector pointing from C_*n*−1_ to N_*n*_. The Y-axis, perpendicular to the X-axis, is defined in such a way that C*α*_*n*_ lies in the half-plane Oxy with *y* > 0. The Z-axis is defined as the vector product *X* × *Y*. The origin of the cube is determined in such a way that N_*n*_ is located at position (6.1Å, 6.6Å, 9.6Å). This choice ensures that all the atoms of the central residue fit in the cube. More details can be found in [27]. This representation is invariant to the global orientation of the structure while preserving information about the atoms and residues relative orientations.

### Selection process on SKEMPI V2.0

In the SKEMPI v2.0 database, the mutations happening in the interface (SUP, COR, RIM), in particular in the core (COR), induce bigger changes in binding affinity than the ones located in the non-interacting surface (SUR) or the interior (INT) of the protein (**Fig. SI 3**). Overall, we observed a tendency for the mutations to be deleterious rather than beneficial. The most impactful single-point mutation is located in the complex 1CHO with ΔΔ*G*_*bind*_ = 8.802 kcal/mol. We restricted our experiments to the entries for which the binding affinity were measured by a reliable experimental method, namely ITC, SPR, FL, or SP, as done in [35]. This first filtering step led to 4,974 entries associated with 255 protein complexes. We retained 4,634 entries from 245 complexes by excluding mutation entries with ambiguous free energy or without energy change. We then focused only on the 3,393 single-mutation entries coming from 222 complexes. After removing duplicated entries (a protein complex with the same mutations), we remained with 2,975 mutations. We finally randomly selected a subset of 2,003 mutations associated with 142 complexes. We call this subset *S2003*.

**Figure SI 1:**
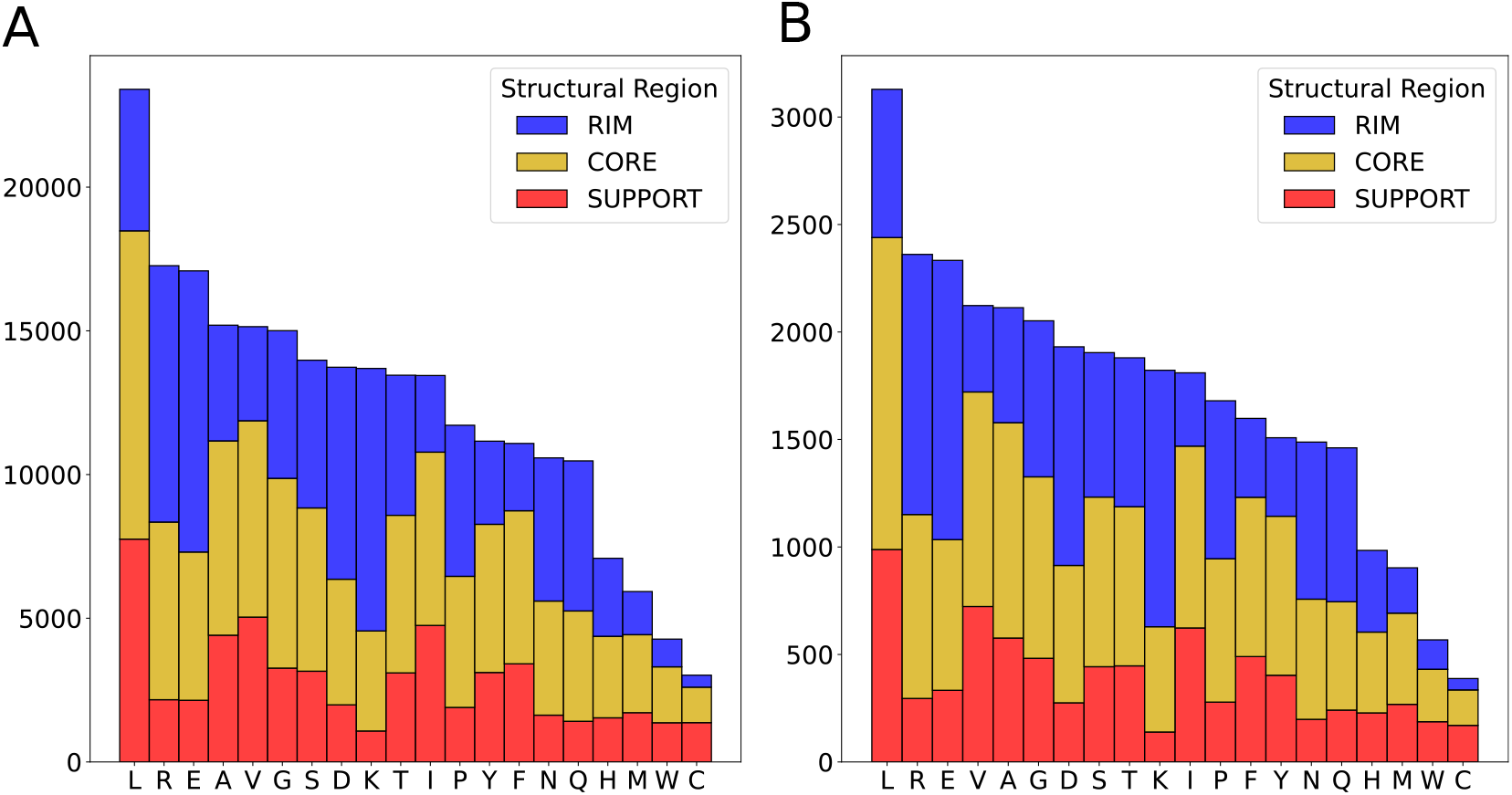
Frequency of interfacial amino acids in the train and validation sets. **A**. Train set. **B**.

**Figure SI 2:**
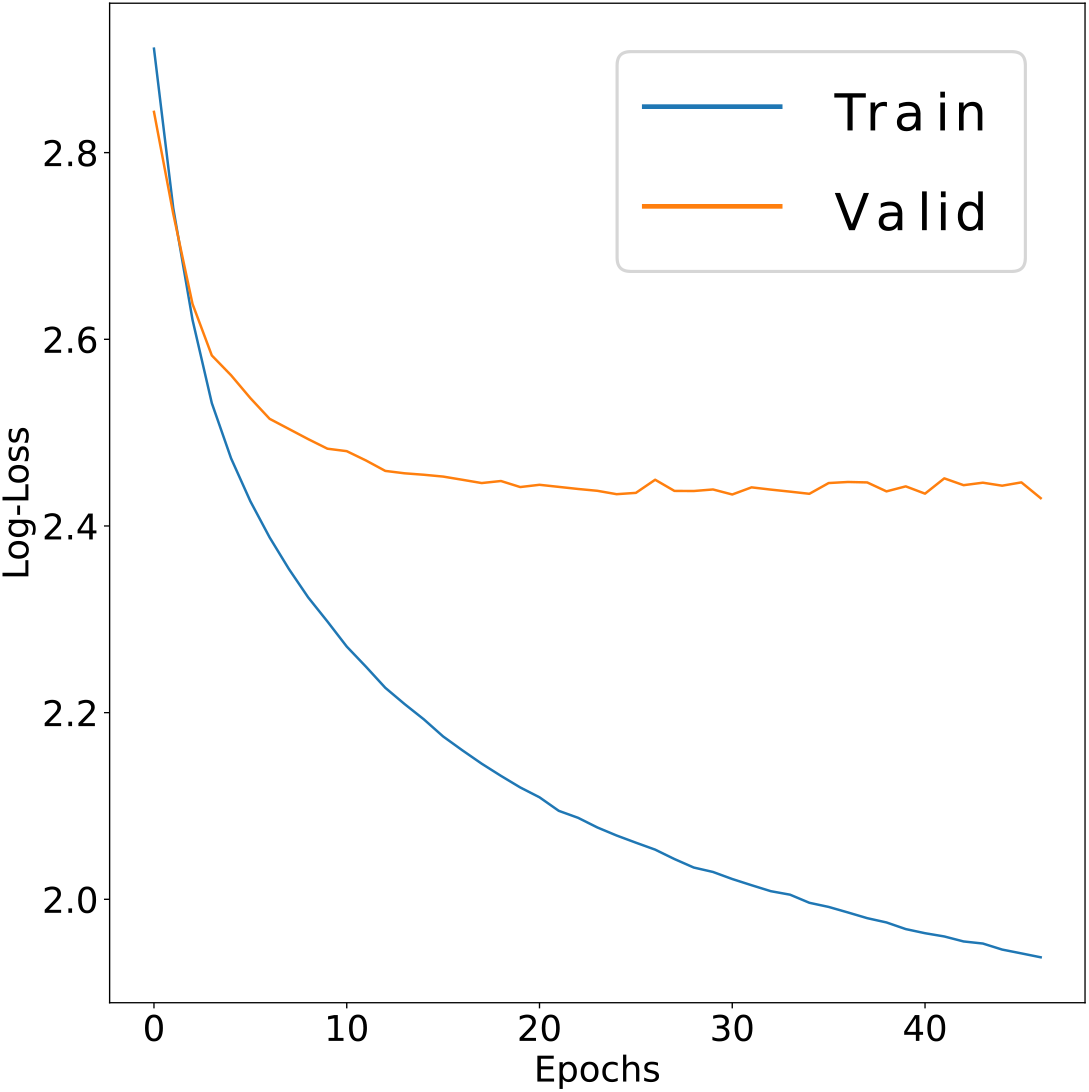
Train and validation loss curves of ssDLA. The x-axis is the number of epochs and the y-axis is the log-loss (categorical cross-entropy). The number of channels is either 167 in case of ssDLA-167 (**A**) or 4 in case of ssDLA-4 (**B**).

### Structural databases

#### Curation of PDBInter

We downloaded all PDB biological assemblies (June 2020 release) from the FTP archive rsync.wwpdb.org::ftp/data/biounit. We discarded the entries with more than 100 chains or with a resolution lower than 5Å. We also removed the protein chains smaller than 20 residues or with more than 20% of unknown residues. We then redundancy-reduced the resulting dataset using annotations from the SCOPe database [14, 7].

#### Backrub modelling and generation of S2003-3D

We generated conformational ensembles for the wild-type and mutated complexes from S2003. We followed a modeling protocol similar to that reported in [2]. It relies on the backrub method [34] for sampling side chain and backbone conformational changes. Our goal was to accurately mimic and explore the fluctuations around a native state. We refer to each generated conformation as a backrub model. We generated 30 backrub models for each wild-type or mutated complex. This amount was shown to be sufficient for estimating free energies in [2].

The protocol unfolds in two optimization steps carried out on the side chains and the backbone (**Fig. SI 4**):

**Figure SI 3:**
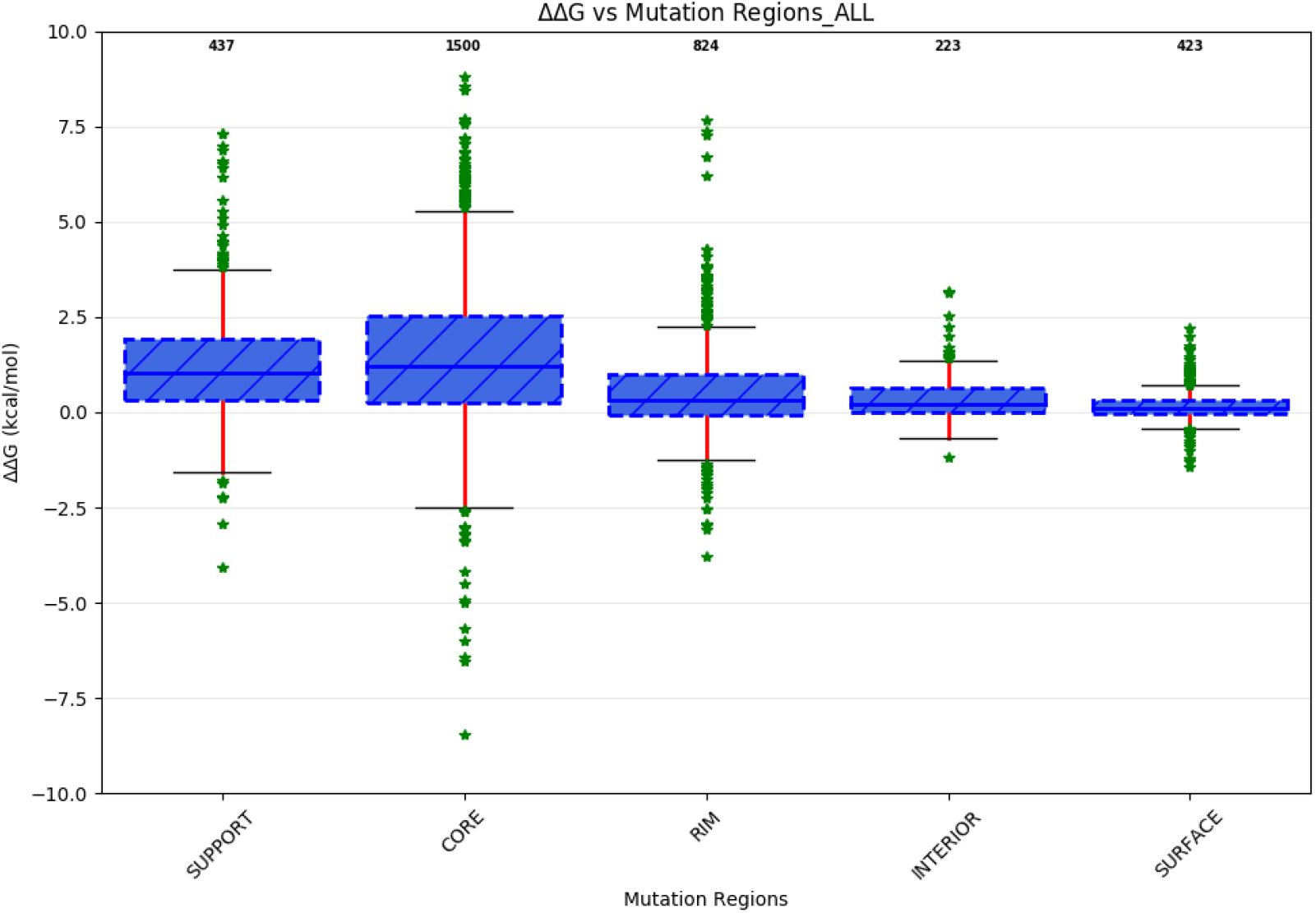
Analysis on SKEMPI V 2.0. ΔΔG associated to single point mutations corresponding to five regions: COR (1500 mutations), SUP (437 mutations), RIM (824 mutations), INT (223 mutations), and SUR (423 mutations).

**Figure SI 4:**
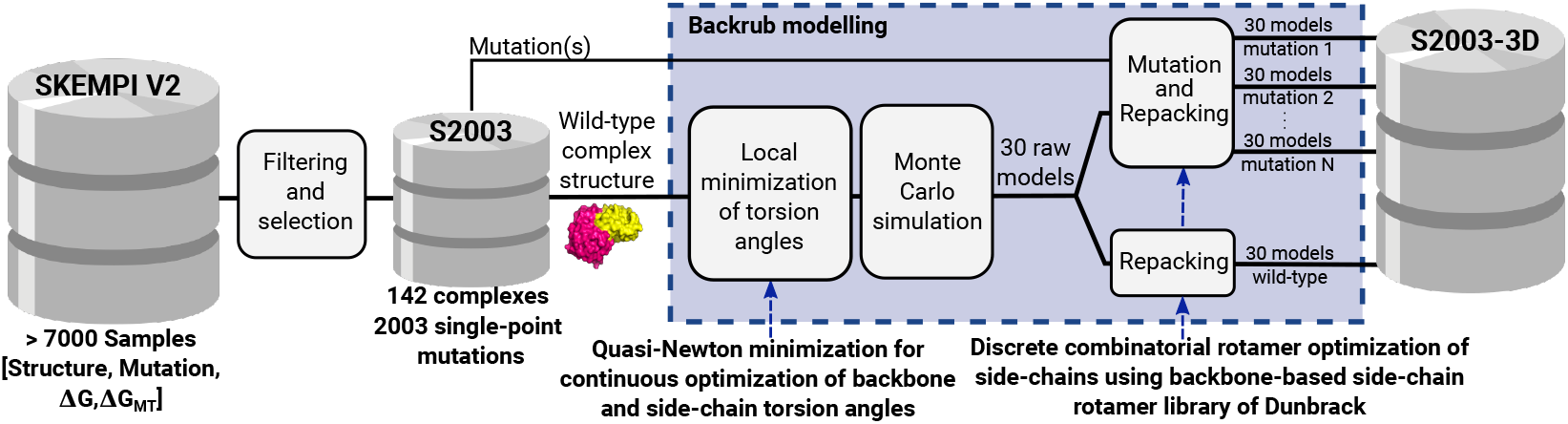
Pipeline for the generation of mutated complexes with backrub. After filtering the SKEMPI V2.0 database, we retained 2003 single point-mutations for 142 complexes (S2003). A wild-type structure undergoes a local minimisation of backbone and side-chain torsion angles followed by a Monte Carlo simulation step. We applied it to produce thirty models for each mutated structure and thirty for the wild-type. This process is followed by a repacking step applied to wild-type and mutation models. For the mutation positions at the interface of each model, we compute the associated cubic volumetric maps.

1. for the backbone and the side chains, it applies quasi-Newton minimization for continuous optimization of torsion angles: Φ, Ψ, *χ*_1_, *χ*_2_, *χ*_3_, etc.
2. for the side chains only, it performs Monte Carlo simulation with the backbone-based side-chain rotamer library of Dunbrack [33] for discrete combinatorial rotamer optimization, also known as repacking.

### Performance assessment

#### Visualisation of ssDLA’s ability to recover the central amino acid

To visualise the performance of the model, we generated logos from pseudo alignments of 20 columns corresponding to the 20 amino acids. In the column corresponding to the amino acid *a*_*i*_, the frequency of occurrence of each amino acid *a*_*j*_ depends on its propensity to be predicted by ssDLA (*i*.*e*. having maximum probability score among the 20 candidate amino acids) when the true central residue of the input cube is a_*i*_. If some amino acid was never predicted, we simply put a gap character. We classify and color the amino acids based on their physico-chemical and geometrical properties (**Table SI 1**). We defined seven classes, namely the aromatic amino acids (ARO: F, W, Y, H), the hydroxyl-containing ones plus Alanine (CAST: C, A, S, T), the aliphatic hydrophobic ones (PHOB: I, L, M, V), the positively charged ones (POS: K, R), the polar and negatively charged ones (POL-N: N, Q, D, E), Glycine (GLY) and Proline (PRO). The classification was taken from [22]. It previously proved relevant for predicting the functional impact of mutations [20].

**Table SI 1:** Seven classes of amino acids.

| Class name | Description | Amino acid(s) | Representative color |
| --- | --- | --- | --- |
| ARO | Aromatic | F, W, Y, H | Green |
| CAST | Hydroxyl-containing and Alanine | C, A, S, T | Black |
| PHOB | Aliphatic hydrophobic | I, L, M, V | Red |
| POS | Positively charged | K, R | Purple |
| POL-N | Polar and negatively charged | N, Q, D, E | Blue |
| GLY | Glycine | G | Gray |
| PRO | Proline | P | Orange |

#### Change of the amino acid size upon mutation

We calculated the change of amino acid size as a volume difference (*δV*) between wild-type and mutant following [16]. A mutation was classified as “neutral” if |*δV* | < 10Å^3^, as small to large if *δV* > 10Å^3^, and as large to small if *δV* < −10Å^3^.

**Figure SI 5:**
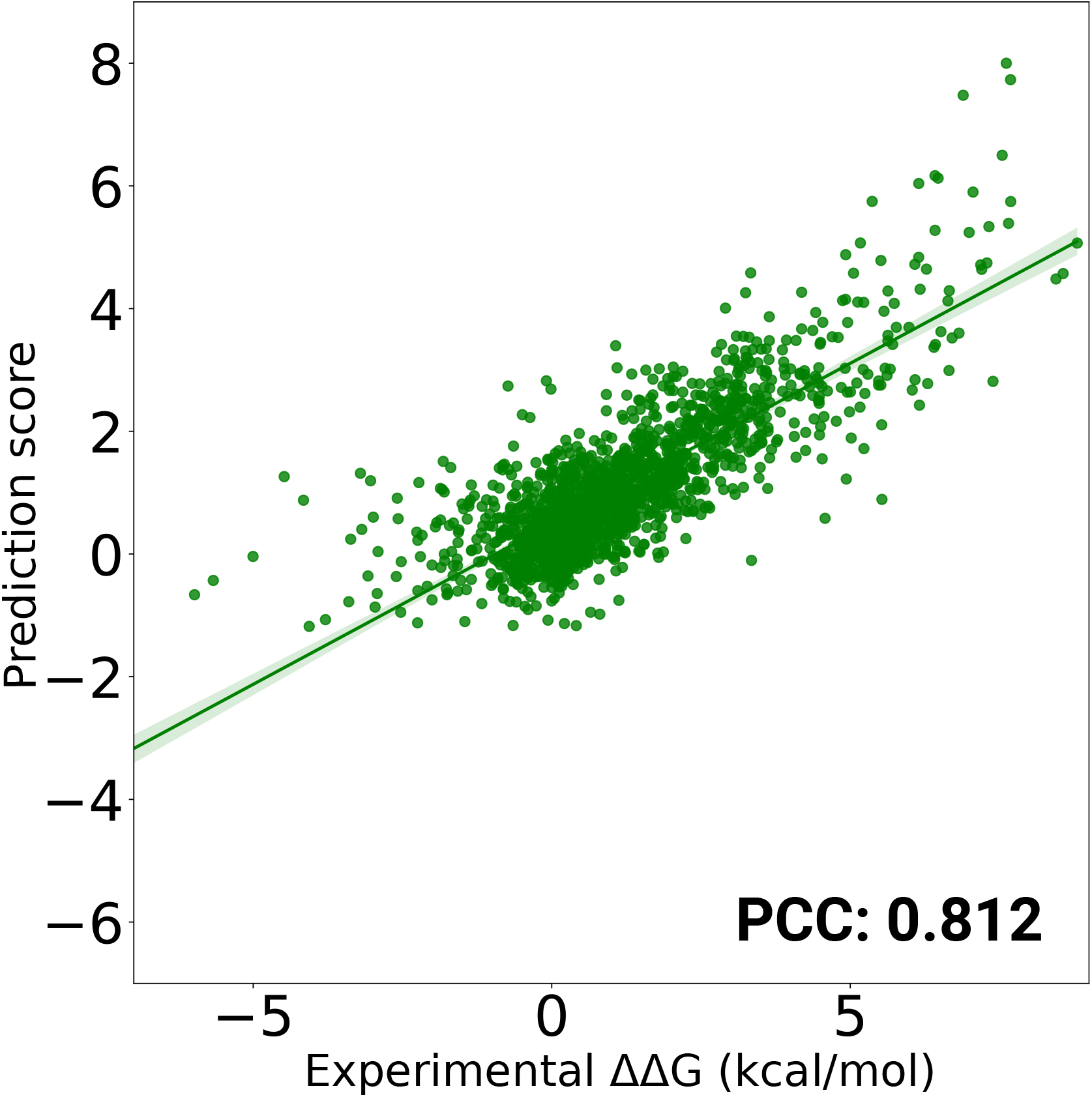
The predictive performance of DLA-Mutation is evaluated on 2003 mutations following a mutation-based 10-fold cross validation. The experimental setup is with pre-training and including complete set of auxiliary features.

## Notes

### Competing Interest Statement

The authors have declared no competing interest.

## References

[1] Namrata Anand, Raphael Eguchi, Irimpan I. Mathews, Carla P. Perez, Alexander Derry, Russ B. Altman, and Po-Ssu Huang. Protein sequence design with a learned potential. Nature Commu-nicationsd, 13(1):746, February 2022.

[2] Kyle A. Barlow, Shane Ó Conchúir, Samuel Thompson, Pooja Suresh, James E. Lucas, Markus Heinonen, and Tanja Kortemme. Flex ddG: Rosetta Ensemble-Based Estimation of Changes in Protein–Protein Binding Affinity upon Mutation. The Journal of Physical Chemistry B, 122(21):5389–5399, May 2018.

[3] Tristan Bepler and Bonnie Berger. Learning the protein language: Evolution, structure, and function. Cell Systems, 12(6):654–669.e3, June 2021.

[4] H. M. Berman, T. Battistuz, T. N. Bhat, W. F. Bluhm, P. E. Bourne, K. Burkhardt, Z. Feng, G. L. Gilliland, L. Iype, S. Jain, P. Fagan, J. Marvin, D. Padilla, V. Ravichandran, B. Schneider, N. Thanki, H. Weissig, J. Westbrook, and C. Zardecki. The Protein Data Bank. Acta Crystallographica Section D: Biological Crystallography, 58(6):899–907, June 2002.

[5] Lasse M. Blaabjerg, Maher M. Kassem, Lydia L. Good, Nicolas Jonsson, Matteo Cagiada, Kristoffer E. Johansson, Wouter Boomsma, Amelie Stein, and Kresten Lindorff-Larsen. Rapid protein stability prediction using deep learning representations, August 2022. Pages: 2022.07.14.500157 Section: New Results.

[6] Nicoletta Ceres, Marco Pasi, and Richard Lavery. A Protein Solvation Model Based on Residue Burial. Journal of Chemical Theory and Computation, 8(6):2141–2144, June 2012. Publisher: American Chemical Society.

[7] John-Marc Chandonia, Lindsey Guan, Shiangyi Lin, Changhua Yu, Naomi K Fox, and Steven E Brenner. SCOPe: improvements to the structural classification of proteins – extended database to facilitate variant interpretation and machine learning. Nucleic Acids Research, 50(D1):D553–D559, January 2022.

[8] J. Dauparas, I. Anishchenko, N. Bennett, H. Bai, R. J. Ragotte, L. F. Milles, B. I. M. Wicky, A. Courbet, R. J. de Haas, N. Bethel, P. J. Y. Leung, T. F. Huddy, S. Pellock, D. Tischer, F. Chan, B. Koepnick, H. Nguyen, A. Kang, B. Sankaran, A. K. Bera, N. P. King, and D. Baker. Robust deep learning based protein sequence design using ProteinMPNN, June 2022. Pages: 2022.06.03.494563 Section: New Results.

[9] Alessia David, Rozami Razali, Mark N. Wass, and Michael J.E. Sternberg. Protein–protein interaction sites are hot spots for disease-associated nonsynonymous SNPs. Human Mutation, 33(2):359–363, 2012. _eprint: https://onlinelibrary.wiley.com/doi/pdf/10.1002/humu.21656.

[10] Alessia David and Michael J. E. Sternberg. The Contribution of Missense Mutations in Core and Rim Residues of Protein–Protein Interfaces to Human Disease. Journal of Molecular Biology, 427(17):2886–2898, August 2015.

[11] Jacob Devlin, Ming-Wei Chang, Kenton Lee, and Kristina Toutanova. BERT: Pre-training of Deep Bidirectional Transformers for Language Understanding. In Proceedings of the 2019 Conference of the North American Chapter of the Association for Computational Linguistics: Human Language Technologies, Volume 1 (Long and Short Papers), pp. 4171–4186, Minneapolis, Minnesota, June 2019. Association for Computational Linguistics.

[12] Ahmed Elnaggar, Michael Heinzinger, Christian Dallago, Ghalia Rehawi, Yu Wang, Llion Jones, Tom Gibbs, Tamas Feher, Christoph Angerer, Martin Steinegger, Debsindhu Bhowmik, and Burkhard Rost. ProtTrans: Towards Cracking the Language of Lifes Code Through Self-Supervised Deep Learning and High Performance Computing. IEEE Transactions on Pattern Analysis and Machine Intelligence, pp. 1–1, 2021. Conference Name: IEEE Transactions on Pattern Analysis and Machine Intelligence.

[13] Stefan Engelen, Ladislas A. Trojan, Sophie Sacquin-Mora, Richard Lavery, and Alessandra Carbone. Joint Evolutionary Trees: A Large-Scale Method To Predict Protein Interfaces Based on Sequence Sampling. PLOS Computational Biology, 5(1):e1000267, January 2009. Publisher: Public Library of Science.

[14] Naomi K. Fox, Steven E. Brenner, and John-Marc Chandonia. SCOPe: Structural Classification of Proteins—extended, integrating SCOP and ASTRAL data and classification of new structures. Nucleic Acids Research, 42(D1):D304–D309, January 2014.

[15] Cunliang Geng, Anna Vangone, Gert E. Folkers, Li C. Xue, and Alexandre M. J. J. Bonvin. iSEE: Interface structure, evolution, and energy-based machine learning predictor of binding affinity changes upon mutations. Proteins: Structure, Function, and Bioinformatics, 87(2):110–119, 2019.

[16] Yehouda Harpaz, Mark Gerstein, and Cyrus Chothia. Volume changes on protein folding. Structure, 2(7):641–649, July 1994.

[17] Chloe Hsu, Robert Verkuil, Jason Liu, Zeming Lin, Brian Hie, Tom Sercu, Adam Lerer, and Alexander Rives. Learning inverse folding from millions of predicted structures. In Proceedings of the 39th International Conference on Machine Learning, pp. 8946–8970. PMLR, June 2022. ISSN: 2640-3498.

[18] S.J. Hubbard and J.M. Thornton. NACCESS, Computer Program, 1993.

[19] Justina Jankauskaitė, Brian Jiménez-García, Justas Dapkūnas, Juan Fernández-Recio, and Iain H. Moal. SKEMPI 2.0: an updated benchmark of changes in protein–protein binding energy, kinetics and thermodynamics upon mutation. Bioinformatics, 35(3):462–469, February 2019.

[20] Elodie Laine, Yasaman Karami, and Alessandra Carbone. GEMME: A Simple and Fast Global Epistatic Model Predicting Mutational Effects. Molecular Biology and Evolution, 36(11):2604–2619, November 2019.

[21] Emmanuel D. Levy. A Simple Definition of Structural Regions in Proteins and Its Use in Analyzing Interface Evolution. Journal of Molecular Biology, 403(4):660–670, November 2010.

[22] Tanping Li, Ke Fan, Jun Wang, and Wei Wang. Reduction of protein sequence complexity by residue grouping. Protein Engineering, Design and Selection, 16(5):323–330, May 2003.

[23] Xianggen Liu, Yunan Luo, Sen Song, and Jian Peng. Pre-training of Graph Neural Network for Modeling Effects of Mutations on Protein-Protein Binding Affinity. arXiv:2008.12473 [cs, q-bio], August 2020. arXiv: 2008.12473.

[24] Mihaly Mezei. A new method for mapping macromolecular topography. Journal of Molecular Graphics and Modelling, 21(5):463–472, March 2003.

[25] Yasser Mohseni Behbahani, Simon Crouzet, Elodie Laine, and Alessandra Carbone. Deep Local Analysis evaluates protein docking conformations with locally oriented cubes. Bioinformatics, p. btac551, August 2022.

[26] Surendra S. Negi and Werner Braun. Statistical analysis of physical-chemical properties and prediction of protein-protein interfaces. Journal of Molecular Modeling, 13(11):1157–1167, November 2007.

[27] Guillaume Pagès, Benoit Charmettant, and Sergei Grudinin. Protein model quality assessment using 3D oriented convolutional neural networks. Bioinformatics (Oxford, England), 35(18):3313–3319, September 2019.

[28] Douglas E. V. Pires, David B. Ascher, and Tom L. Blundell. mCSM: predicting the effects of mutations in proteins using graph-based signatures. Bioinformatics (Oxford, England), 30(3):335–342, February 2014.

[29] Douglas E.V. Pires and David B. Ascher. mCSM-AB: a web server for predicting anti-body–antigen affinity changes upon mutation with graph-based signatures. Nucleic Acids Research, 44(W1):W469–W473, July 2016.

[30] Alexander Rives, Joshua Meier, Tom Sercu, Siddharth Goyal, Zeming Lin, Jason Liu, Demi Guo, Myle Ott, C. Lawrence Zitnick, Jerry Ma, and Rob Fergus. Biological structure and function emerge from scaling unsupervised learning to 250 million protein sequences. Proceedings of the National Academy of Sciences, 118(15), April 2021. Publisher: National Academy of Sciences Section: Biological Sciences.

[31] Carlos H. M. Rodrigues, Yoochan Myung, Douglas E. V. Pires, and David B. Ascher. mCSM-PPI2: predicting the effects of mutations on protein–protein interactions. Nucleic Acids Research, 47(W1):W338–W344, July 2019. Publisher: Oxford Academic.

[32] Carlos H M Rodrigues, Douglas E V Pires, and David B Ascher. mmCSM-PPI: predicting the effects of multiple point mutations on protein–protein interactions. Nucleic Acids Research, 49(W1):W417–W424, July 2021.

[33] Maxim V. Shapovalov and Roland L. Dunbrack. A Smoothed Backbone-Dependent Rotamer Library for Proteins Derived from Adaptive Kernel Density Estimates and Regressions. Structure, 19(6):844–858, June 2011.

[34] Colin A. Smith and Tanja Kortemme. Backrub-Like Backbone Simulation Recapitulates Natural Protein Conformational Variability and Improves Mutant Side-Chain Prediction. Journal of Molecular Biology, 380(4):742–756, July 2008.

[35] Anna Vangone and Alexandre MJJ Bonvin. Contacts-based prediction of binding affinity in protein–protein complexes. eLife, 4:e07454, July 2015. Publisher: eLife Sciences Publications, Ltd.

[36] Menglun Wang, Zixuan Cang, and Guo-Wei Wei. A topology-based network tree for the prediction of protein–protein binding affinity changes following mutation. Nature Machine Intelligence, 2(2):116–123, February 2020. Number: 2 Publisher: Nature Publishing Group.

[37] Dapeng Xiong, Dongjin Lee, L. Li, Qiuye Zhao, and Haiyuan Yu. Implications of disease-related mutations at protein–protein interfaces. Current Opinion in Structural Biology, 72:219–225, February 2022.

[38] Peng Xiong, Chengxin Zhang, Wei Zheng, and Yang Zhang. BindProfX: Assessing Mutation-Induced Binding Affinity Change by Protein Interface Profiles with Pseudo-Counts. Journal of Molecular Biology, 429(3):426–434, February 2017.

[39] Zuobai Zhang, Minghao Xu, Arian Jamasb, Vijil Chenthamarakshan, Aurelie Lozano, Payel Das, and Jian Tang. Protein Representation Learning by Geometric Structure Pretraining, May 2022. arXiv:2203.06125 [cs].

[40] Guangyu Zhou, Muhao Chen, Chelsea J T Ju, Zheng Wang, Jyun-Yu Jiang, and Wei Wang. Mutation effect estimation on protein–protein interactions using deep contextualized representation learning. NAR Genomics and Bioinformatics, 2(2):lqaa015, June 2020.

